# Systems-level identification of transcription factors critical for mouse embryonic development

**DOI:** 10.1101/167197

**Authors:** Kai Zhang, Mengchi Wang, Ying Zhao, Wei Wang

## Abstract

Dynamic changes in the transcriptional regulatory circuit can influence the specification of distinct cell types. Numerous transcription factors (TFs) have been shown to function through dynamic rewiring during embryonic development but a comprehensive survey on the global regulatory network is still lacking. Here, we performed an integrated analysis of epigenomic and transcriptomic data to reveal key regulators from 2 cells to postnatal day 0 in mouse embryogenesis. We predicted 3D chromatin interactions including enhancer-promoter interactions in 12 tissues across 8 developmental stages, which facilitates linking TFs to their target genes for constructing genetic networks. To identify driver TFs particularly those not necessarily differentially expressed ones, we developed a new algorithm, dubbed as Taiji, to assess the global importance of TFs in development. Through comparative analysis across tissues and developmental stages, we systematically uncovered TFs that are critical for lineage-specific and stage-dependent tissue specification. Most interestingly, we have identified TF combinations that function in spatiotemporal order to form transcriptional waves regulating developmental progress and differentiation. Not only does our analysis provide the first comprehensive map of transcriptional regulatory circuits during mouse embryonic development, the identified novel regulators and the predicted 3D chromatin interactions also provide a valuable resource to guide further mechanistic studies.

## Introduction

Transcription factors (TFs) are important regulators of cell fate and play pivotal roles in development^1^. Identification of driver TFs at different developmental stages would help to reveal how tissue differentiation is achieved and shed light on understanding the regulatory mechanisms. The dynamic activities of TFs are partially due to the temporal variation of their own transcription levels. Effort of identifying driver TFs has been thus largely focused on differentially expressed genes (DEGs) across tissues and/or time points. However, this approach is problematic as TFs’ activity is not necessarily correlated with their expression level^2^. Therefore it would miss regulators that themselves do not present significant expression variations but instead regulate many DEGs or other critical TFs.

For the first time, the ENCODE project has systematically mapped the epigenomic dynamics during mouse embryonic development in 12 tissues and 8 developmental stages. These data provide an unprecedented opportunity to uncover the mechanisms of the developmental regulation but also pose a great challenge for computational analysis. To tackle this challenge, we have developed a framework, dubbed as Taiji, to incorporate different genomic information, including chromatin state (from chromatin immunoprecipitation followed by high-throughput sequencing (ChIP-seq) or assay for transposase-accessible chromatin with high throughput sequencing (ATAC-seq)), gene expression profile (from RNA sequencing (RNA-seq) or microarray), and chromatin long range interactions (from Hi-C or computational prediction), to construct genetic networks. By leveraging novel graph algorithms, we have identified key TFs crucial in defining tissue differentiation. Furthermore, we applied Taiji to the existing data in earlier development, from 2-cell stage to embryonic stem cell, which complements the data generated by the mouse ENCODE project. Our analyses thus provide the first comprehensive identification of key regulators for a variety of tissues during mouse embryogenesis. While many of the identified regulators are well supported by the literature, the newly discovered key TFs of development provide a valuable resource for follow up biological analysis. Most interestingly, we uncovered TF combinations that are activated in a spatiotemporal manner, which behave like transcriptional waves to direct the developmental progress and tissue differentiation.

## Results

### An overview of the Taiji framework

Taiji integrates diverse data sets to identify key regulators, including genome-wide expression profile, chromatin state and long-range chromatin interactions. The software has a user-friendly command line interface which is implemented in Haskell. Taiji can be easily set up and run in desktop computers. Furthermore, Taiji is designed to handle the big data through tightly integrated with workload managers that support the distributed resource management application API (DRMAA), such as the open grid scheduler and the slurm workload manager. This feature allows Taiji to analyze numerous data sets in parallel on high performance computing clusters. In addition, Taiji has a built-in workflow manager, featuring automatic checkpointing and data recovery, allowing continuation of the analysis after interruption by error crash or power outage.

Taiji builds a genetic network by assembling regulatory interactions between TFs and genes. Based on this network, the personalized PageRank algorithm^3^ is used to identify key regulators that are associated with DEGs (Fig. 1a and Extended Data Fig. 1). Taiji starts with identifying active regulatory elements, including active promoters and active enhancers, defined by ATAC-seq or H3K27ac ChIP-seq peaks. To link enhancers to their target genes, Taiji provide several options. Under the default settings assuming no 3D genome information is available, Taiji assigns enhancers to the nearest genes. The limitation of such approach is obvious and it may miss many long-range enhancer-promoter interactions. Although Taiji can take advantage of Hi-C experiments to identify spatially associated enhancer-gene pairs, Hi-C is still expensive and requires a large amount of materials that is not available in this project. To overcome these limitations, here we applied an improved version of our previously developed computational method, EpiTensor^4^, to predict 3D chromatin interactions and assign enhancers to their interacting promoters. Next, Taiji scans known TF motifs^5^ within each regulatory element to predict their putative binding sites. TFs with putative binding sites in promoters or enhancers are then linked to their target genes to form a network.

**Figure 1:**
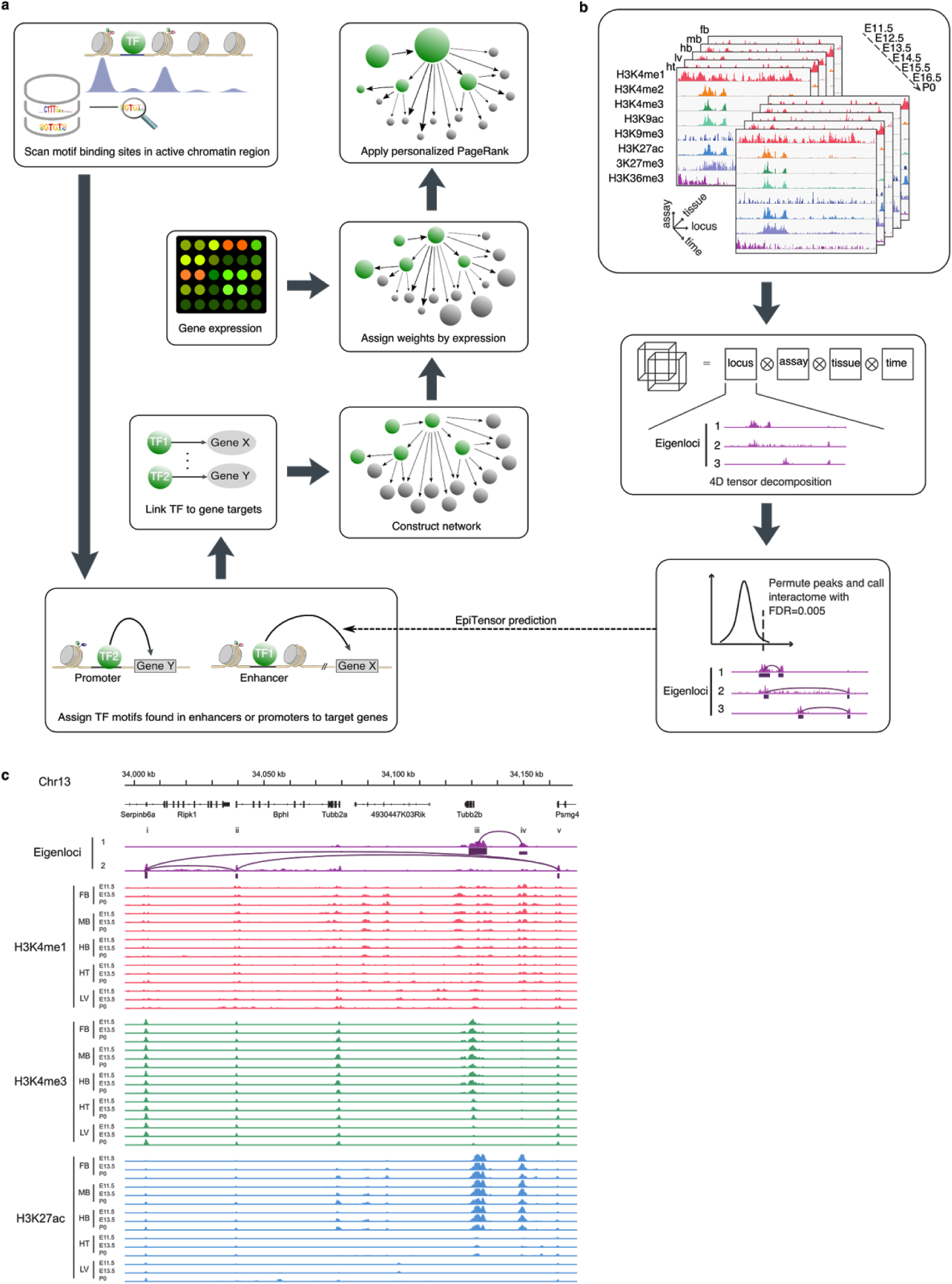
The design of Taiji and EpiTensor algorithms. **a**, The workflow of Taiji pipeline. **b**, The workflow of EpiTensor algorithm. **c**, An example of chromatin interactions predicted by EpiTensor. Locus iv is a validated enhancer documented in the Vista database.

A TF’s activity is not necessarily correlated with its expression level^2^. We argue that the expression variations of TFs’ regulatees is a better indicator of their activities. For this purpose, we applied the personalized PageRank algorithm to ranking TFs in a genetic network. In our formulation, a TF’s rank is initially determined by the number of DEGs among its regulatees. Afterwards, the weights received from its regulatees are propagated to its parent nodes. As a result, a TF is ranked high if it regulates many DEGs or it is linked to other highly ranked TFs or both. Mathematically, the personalized PageRank is the stationary distribution of a random walk which, at each step, with a certain probability *α* jumps to a random node, and with probability 1 – *α* follows a randomly chosen outgoing edge from the current node. Formally, let *G* = (*V, E*) denote a directed graph, where *V* is a set of nodes and *E* contains a directed edge 〈*u, v*〉 if and only if node *u* links to node *v*. Let *A* be the transition matrix. We define 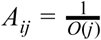 if node *j* links to node *i*, and *A_ij_* = 0 otherwise, where *O*(*j*) is the out-degree of node *j*. Given a seed vector *s*, the personalized PageRank vector *v* is calculated by *v* = (1 – *α*) *Av* + *αs*. In the regulatory network, we set the weight of each gene to 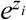, where *z_i_* is the z-score of expression levels of gene *i* in different cell states. The weights of genes are then normalized and used as the seed vector *s* for computing personalized PageRank.

### EpiTensor predicts long range chromatin interactions

One of Taiji’s major innovations is to leverage the computationally predicted chromatin interactome to construct networks. For this purpose, we used our previously published method EpiTensor. EpiTensor provides an unsupervised learning method to predict chromatin interactome at a high resolution of 200 bp within topologically associating domains (TADs) from histone modification data (Fig. 1b). All the predicted 3D interactions are available at http://wanalab.ucsd.edu/star/MouseENCODE/driverTF/index.html. Previously, we have demonstrated that EpiTensor outperforms other correlation-based methods and nearest-gene assignment^4^. EpiTensor successfully identified interactions between many validated enhancers and genes important for development. The predicted interactions cover 25,358 TSS, >38% of all annotated transcripts in Gencode (vM14)^6^ and 334 experimentally validated enhancers in the Vista database^7^, 55% of all confirmed active enhancers during embryonic mouse development.

An example of predicted interactions is shown in Figure 1c (only H3K4me1, H3K4me3 and H3K27ac in E11.5, E13.5 and P0 are shown in the figure due to limited space). Locus i, ii and v are three active promoters marked by H3K4me3 and H3K27ac in the tissue-stages shown in the figure. Locus iii is the promoter of TUBB2B, preferentially active in brain, which is consistent with the reported role of TUBB2B in cortical formation and brain morphogenesis^8^. Locus iv is an enhancer marked by strong H3K4me1 and H3K27ac signals in the brain tissues, and it is an confirmed enhancer active in embryonic mouse midbrain, hindbrain and facial mesenchyme in the Vista database (peak ID: mm1605). Eigenlocus 1 contains a strong peak representing the interaction between TUBB2B promoter at locus iii and the confirmed enhancer at locus iv; the co-variation of histone modifications at the two loci can be exemplified by the coherent decrease of H3K4me1/H3K4me3 and H3K27ac at locus iii and iv from brain to other tissues. Interestingly, the interaction between locus iv and v was not found in any eigenlocus due to lack of such co-variation of the histone marks, despite their closer distance. Note that the distance of locus i and v is large (160 kbp), which indicates the power of EpiTensor for identifying long-range interactions.

### Taiji reveals driver TFs during embryogenesis

Using the predicted promoter-enhancer interactions from EpiTensor, we next constructed TF-gene regulatory networks in 12 tissues from 8 developmental stages. The average number of nodes and edges in these networks are 16,187 and 320,227 respectively. 3.95% (639) of the nodes are TFs. On average, each TF regulates 501 genes, and each gene is regulated by 19 TFs (Extended Data Fig. 2). We then used the PageRank algorithm to rank TFs in individual network. To reveal the differential activities of TFs between tissues or stages, we filtered out the TFs that present less activity variations (coefficients of variance of ranking scores across tissues and stages is less than 1). In Fig. 2a, we showed the ranking scores of 245 most variable TFs. We found that different tissues have distinct TF regulatory patterns, highlighting the important roles of transcriptional regulation in mouse embryonic development. In contrast, we did not discover any discernible patterns for individual TFs across different stages (Fig. 2b). To investigate to what extend the tissue specificity is determined at the transcriptional regulation level, we constructed the lineage tree of 12 tissue from 8 stages using the 245 TFs’ ranking scores as the input. Remarkably, TF ranks alone achieved great accuracy in distinguishing different tissues (Fig. 2c). Except that forebrain, midbrain and craniofacial cannot be separated at the very early stage (E10.5 or E11.5), all other tissues were correctly segregated. As a comparison, a lineage tree constructed using the RNA expression profile of the same 245 TFs results in more errors (Fig. 2d). This further demonstrates that gene expression cannot accurately reflect TF’s transcriptional activity and our metric is superior for this purpose.

**Figure 2:**
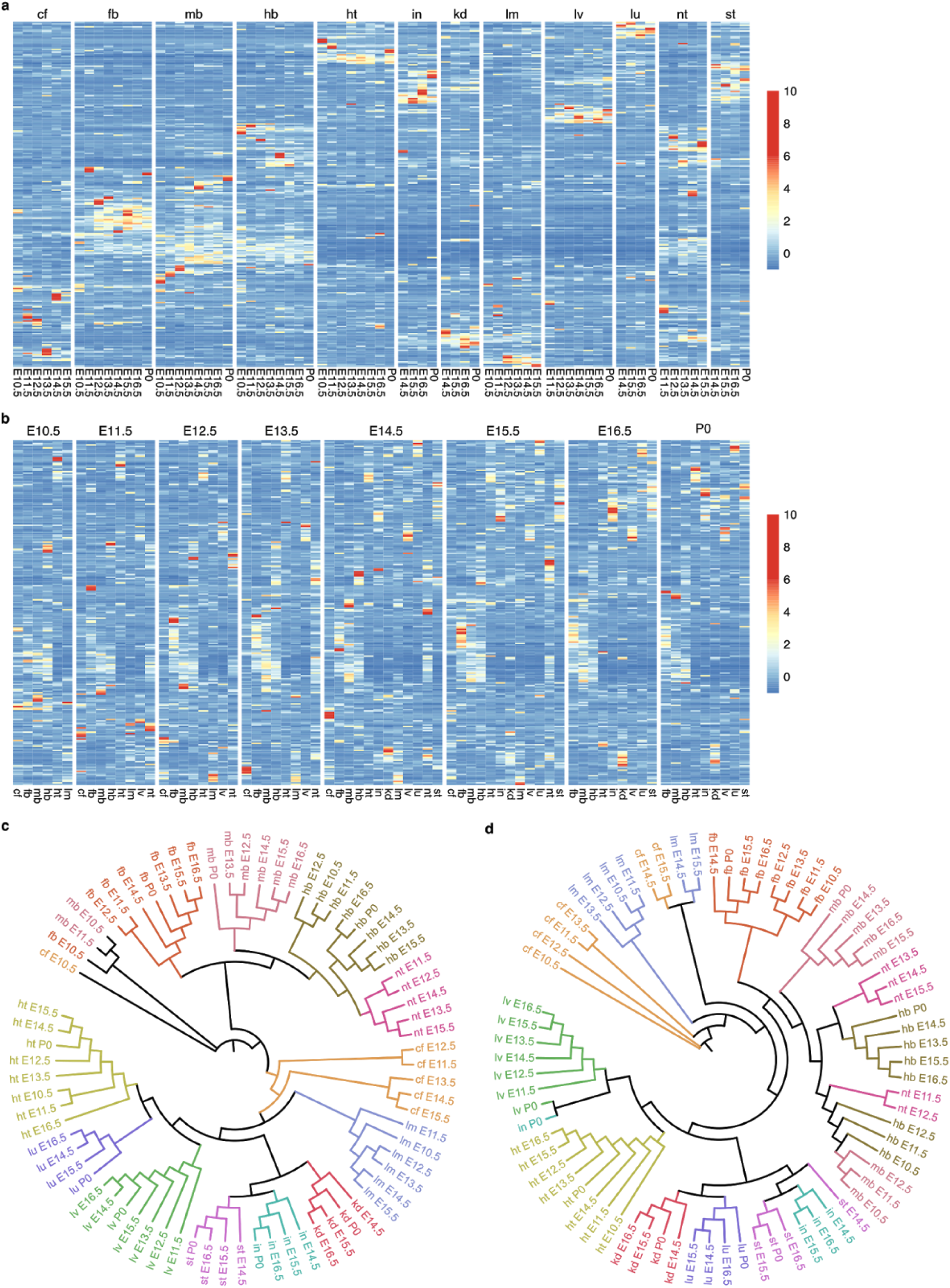
TF activity determined from Taiji accurately predicts tissue specification. Two different views, arranged by tissue types **(a)** and arranged by stages **(b)**, of the 245 most variable TFs’ ranking scores during embryogenesis. **c**, A lineage tree constructed from the ranking scores of the 245 TFs. **d**, A lineage tree constructed from the gene expression levels, determined by RNA-seq, of the same 245 TFs in **(c)**. cf, craniofacial prominence; fb, forebrain; hb, hindbrain; ht, heart; in, intestine; kd, kidney; lm, limb; lv, liver; lu, lung; mb, midbrain; nt, neural tube; st, stomach.

The earliest developmental stage that mouse ENCODE project has surveyed starts from E10.5. To obtain a more complete view of mouse embryogenesis, we incorporated another published data set which profiled the transcriptome and genome-wide chromatin accessibility in earlier mouse embryos^10^, spanning stages from E1 (2-cell) to E5 (blastocyst stage). Based on the analysis of TFs’ ranking scores in different tissues and at various stages (see Methods for details), 35 housekeeping TFs that exhibit high ranking scores across all tissues and stages, were identified (Fig. 3a). Functional classification of these TFs revealed that metabolic process, cellular process and developmental process are the top 3 enriched GO terms, highlighting their roles in basic cellular functions and embryonic development. More interestingly, we have also identified potential driver TFs in 12 tissues from 8 stages, as well as those in 2-cell, 4-cell, 8-cell embryos, inner cell masses (ICMs) and mouse embryonic stem cells (mESCs). For the first time, the complex transcriptional regulation during mouse embryonic development is systematically mapped (Extended Data Fig. 3 and 4).

**Figure 3:**
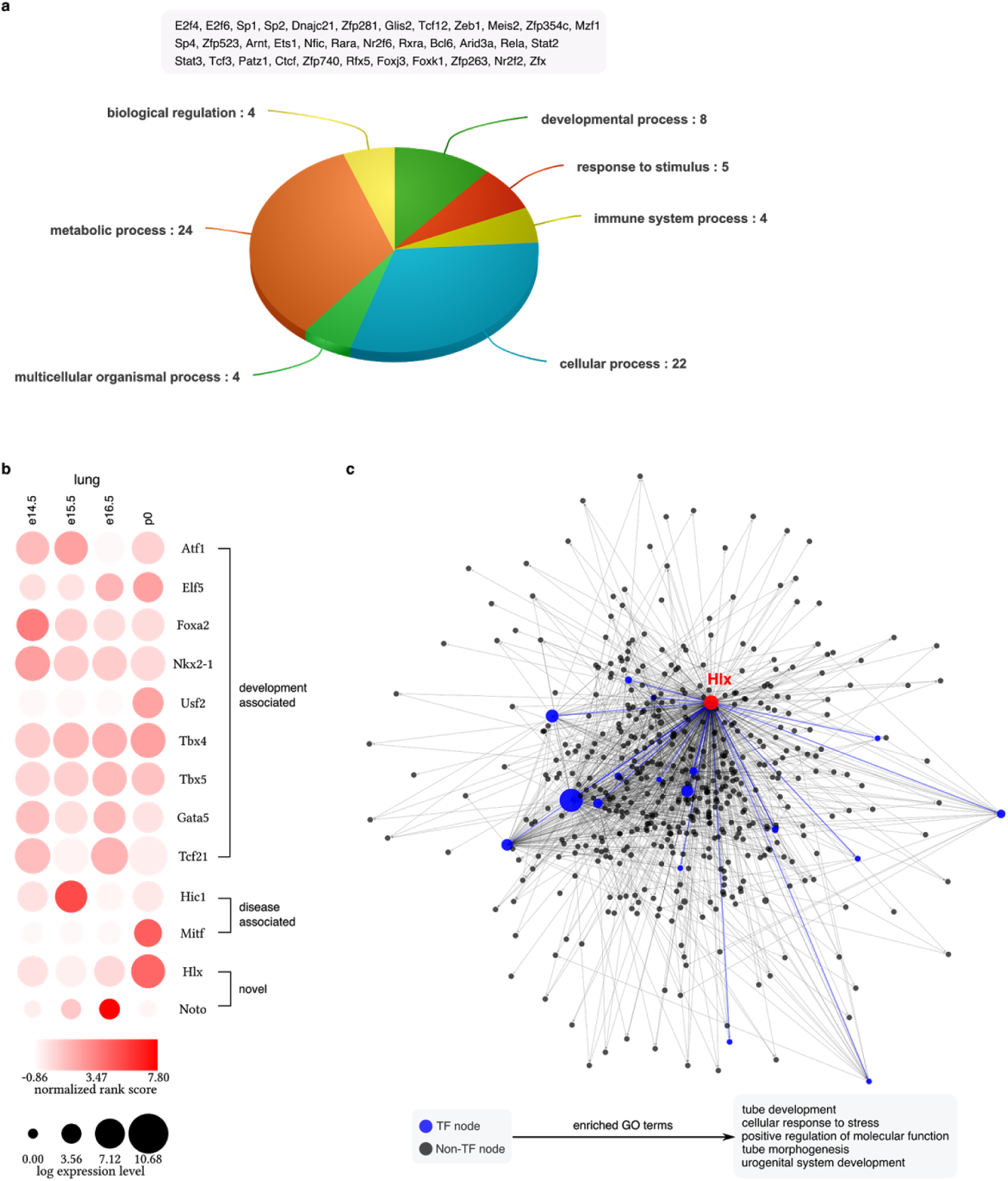
Identification of driver TFs during mouse embryogenesis. **a**, 35 housekeeping TFs that are constitutively active at all stages and in all tissues. Functional classification of these genes reveals their functions in a wide range of biological processes, **b**, Identified driver TFs in lung, including 9 TFs related with lung development, 2 TFs related with lung diseases and 2 novel TFs. **c**, The network for Hlx, a novel driver TF for lung, and its regulatees in lung at p0. Larger nodes represent TFs with higher rank. The bottom shows the top 5 enriched GO terms for Hlx’s regulatees.

The identified driver TFs include many well-known key regulators. For example, we successfully identified *Sox2, Nanog* and *Pou5f1* (also known as *Oct4*) as key TFs in mESCs. To systematically validate our predictions, we performed literature search in 5 tissues that have been extensively studied, including heart, lung, liver, kidney and limb. Remarkably, 41 out of 56 (73.2%) TFs identified by our method are shown to be associated with either development or disease of these tissues (see Supplementary Discussion 1). Taken the lung tissue as an example, 9 out of 13 TFs from our predictions have been previously reported to play a pivotal role during the bronchiole tree and terminal alveolar region formation of mouse lung (Fig. 3b). And another two relate with lung cancer^11,12^. Apart from the spatial specificity (across tissues), temporal pattern of TFs’ activity (across stages) can also be accurately captured by our approach. For instances, *Nkx2-1* is initiated at the early stage of lung development and *Nkx2-1* mutant embryos are arrested at early pseudoglandular (E11-E15) stage^13^, which is consistent with our analysis. Similarly, another key regulator Foxa2, is present in the epithelial cells from the beginning of lung bud formation, and transgenic mice with *Foxa2* ectopically expressed in the lung epithelial cells exhibited defects in branching morphogenesis^14^.

We have also found many novel TFs. *Hlx*, identified as a key regulator for lung development, has not yet been studied in lung. We showed that Hlx is regulating a number of high-rank TFs at P0, including several aforementioned housekeeping TFs, i.e., *Sp1, Meis2* and *Zfx* (Fig. 3c). The functional enrichment analysis of all *Hlx*’s regulatees shows that they likely participate in epithelial tubes development in lung. Besides, some of the predicted novel TFs have been studied in closely related tissues. For instances, *Vsx1*, predicted as an important regulator in hindbrain, was related to retina development^15^. Taken together, these insights revealed by our analysis can guide future functional studies of the developmental mechanisms.

In development, ESCs differentiate into three germ layers (ectoderm, endoderm and mesoderm) and we have identified TFs that are specific to each layer (Fig. 4). For this purpose, tissues originated from the same layer were grouped together and compared against tissues from other layers. Student t-tests with a p-value cutoff of 0.001 were used to identify TFs whose ranks in one germ layer significantly deviate from their ranks in other layers. The functional relevance of the found layer-specific TFs is supported by the literature. For example, we have identified *Zic1, Zic4* and *Zic5* as specific regulators for ectoderm, which is in agreement with the role of *Zic* family in neural development^16^. In additional, many previously known layer-specific markers were also found from our analysis, e.g., *Pax6* and *Otx2* for ectoderm^17,18^, *Foxa2* for endoderm^19^. To further narrow down the aforementioned driver TFs and analyze their tissue specificity, we followed the same strategy to identify tissue-specific TFs within a germ layer by comparing one tissue with other tissues derived from the same layer.

**Figure 4:**
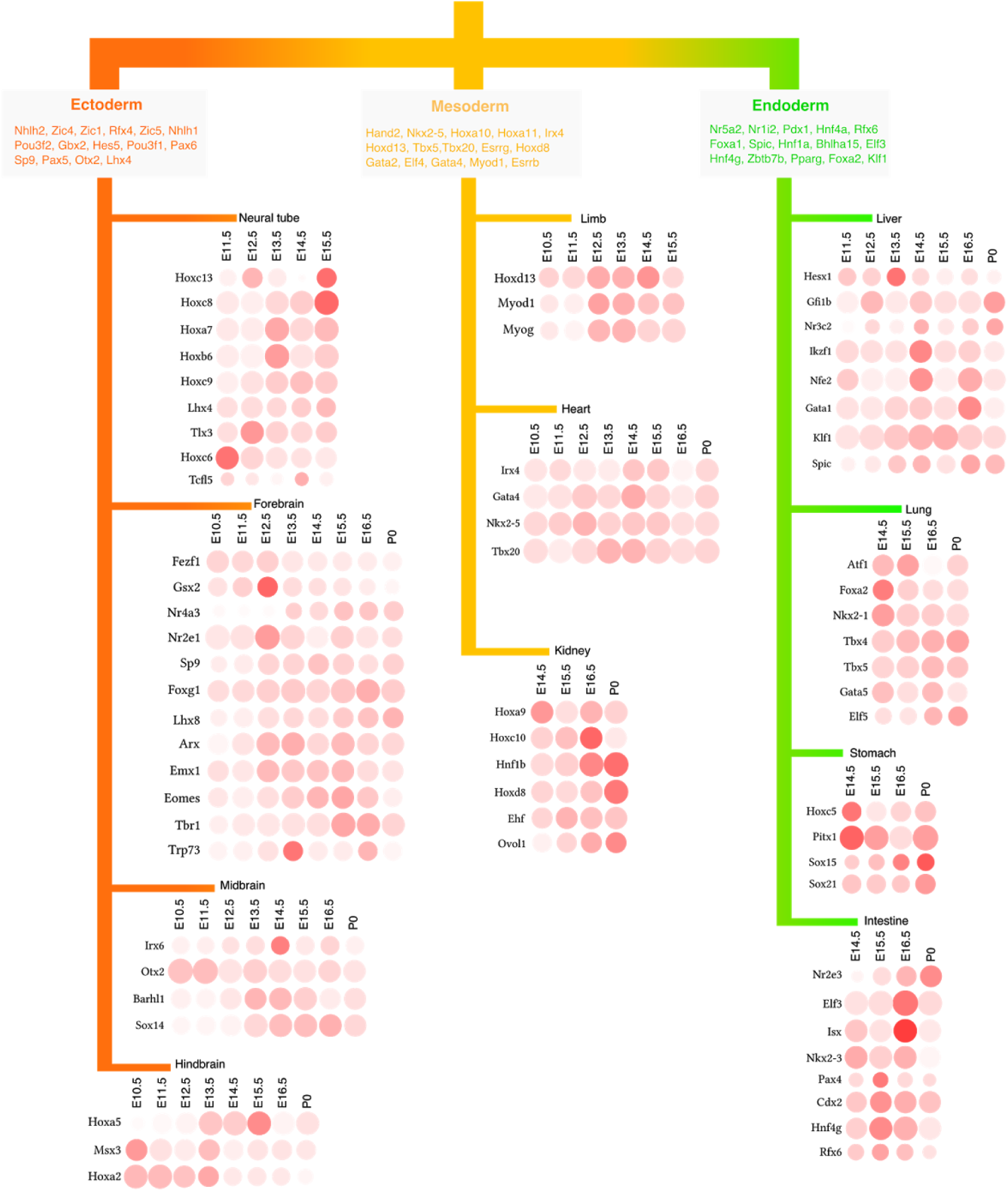
Germ-layer-specific and tissue-specific driver TFs in mouse embryonic development.

### Transcriptional waves during embryogenesis

In addition to the tissue specificity, we have analyzed the temporal activity of TFs during the mouse embryogenesis. To denoise, we applied the principal component analysis (PCA) to the TF ranking score matrix and the scores were then projected to the first 26 out of 77 principal components, corresponding to 70% of the total variance. The projection values were used for subsequent hierarchical clustering analysis. The number of clusters was determined using the dynamic tree cut algorithm^20^. In total, we have identified 44 clusters of TFs that show distinct dynamic patterns during embryogenesis, representing transcriptional waves that orchestrate the tissues differentiation (Extended Data Fig. 5).

Starting from the very early 2-cell stage, waves of transcriptional programs are sequentially turned on. Both C25 (Fig. 4a) and C37 (Extended Data Fig. 5) are initiated at 2-cell stage and TFs in these 2 clusters show highest activity at this stage (or possibly earlier as we do not have data in zygote). However, C25 appears to be more specific to 2-cell stage, as its activity quickly diminishes afterwards. Example TFs from C25 include germ cell specific factor like *Obox1* and *Nr6a1*, both of which are essential for embryogenesis^21,22^, highlighting the roles of parental control in early development. As C25 wanes, C14 and C39 are turned on at 4-cell and 8-cell stage respectively. Subsequently, in ICM and ESC, several waves emerge, including C35, C44, C32, C12. However, these 4 clusters have very distinct dynamics. C35 has a very specific role in ICM and ESC. Well-known pluripotency regulators such as *Pou5f1* and *Nanog* belong to C35. On the contrary, C44 and C32 are more versatile and also play roles in later stages (Fig. 4b). They are also responsible for the differentiation of tissues from specific layers. For example, C44 is highly active in ESC and remains active at later stages in all 4 brain tissues, suggesting its critical role in the regulation of pluripotency and neural differentiation. Indeed, *Sox2* from C44 has been demonstrated to be a pluripotency factor and play a pivotal role in neural differentiation^23^. Similar to C44, C6 also exhibits ectoderm-specific pattern from E10.5 to P0. An important regulator for neural development *Zic5*^16^ belongs to this cluster. In contrast to C44, C32 and C12 are more important in mesoderm and endoderm derived tissues.

Besides germ-layer-specific transcriptional waves, there are also tissue-specific transcriptional programs that drive the differentiation of individual tissue. In Fig. 4c, we highlighted 3 clusters, C1, C29 and C5, which are responsible for stomach/intestine, heart and forebrain development, respectively. Note that we have found such a program for every tissue, including liver (C2), craniofacial (C4, C22, C34), lung (C7), limb (C8), kidney (C15, C33), midbrain (C19, C20), C10 and neural tube/hindbrain (C10). See Extended Data Fig. 5 for details.

Intriguingly, some tissue-specific clusters also appear to have very specific roles at certain stages. Fig. 5d shows that 3 sequential transcriptional waves are responsible for craniofacial development at different stages (C4 for E11.5, C22 for E13.5, C34 for E14.5). As the mechanism of craniofacial development remains largely unknown, our discovery therefore provides an invaluable resource to guide mechanistic studies.

**Figure 5:**
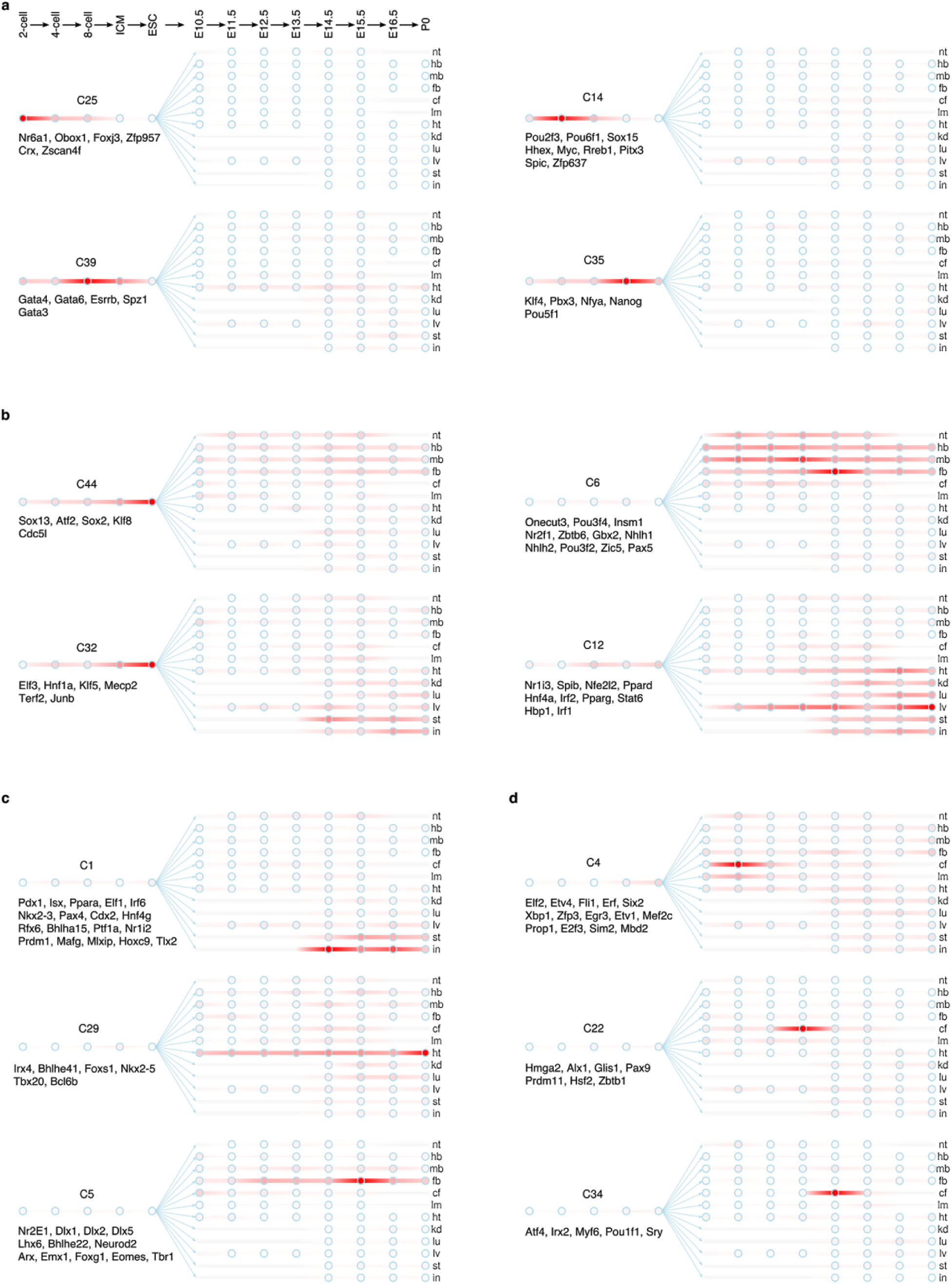
Temporal transcriptional waves direct tissue differentiation during embryogenesis. Circles in each panel represent developmental stages. From left to right, they are 2-cell, 4-cell, 8-cell, ICM, ESC, E10.5, E11.5, E12.5, E13.5, E14.5, E15.5, E16.5 and P0 respectively. TF members are shown below the names of clusters, **a**, 4 waves in early embryonic development. **b**, Layer specific transcriptional waves, **c**, 3 examples of tissue specific transcriptional waves, **d**, Stage-specific waves in craniofacial development.

## Discussion

Here we present a novel and general framework for identifying driver TFs in any biological process. Our method Taiji is capable of flexibly integrating diverse genomic and epigenomic data, including ChIP-seq, RNA-seq, ATAC-seq and Hi-C. Considering the current limitation of Hi-C experiments, we provide a computational alternative when Hi-C experiment is infeasible or unavailable. In this work, we predicted 3D chromatin interactions in 12 tissues and 8 developmental stages of mouse embryo, which provides the first 3D chromatin organization information. By leveraging the strength of diverse experiments, we have successfully mapped lineage-, tissue- and stage-specific driver TFs throughout the mouse embryonic development, from as early as 2-cell stage to postnatal day 0. In addition to retrieving known key regulators, we have also identified new TFs responsible for tissue differentiation and/or development progress. Particularly interesting is the observation of transcriptional waves of TF combinations that are active at particular developmental stages or tissues in temporal order, which can guide the future experimental investigations to understand the regulatory mechanisms of development.

## Methods

### Prediction of chromatin interactions using EpiTensor

TADs were taken from the previous study^24^. The input histone modification data were downloaded from the ENCODE data portal, including a common set of 8 histone marks at 7 time points in 5 tissues: H3K27ac, H3K27me3, H3K36me3, H3K4me1, H3K4me2, H3K4me3, H3K9ac and H3K9me3, at developmental stages of E11.5, E12.5, E13.5, E14.5, E15.5, E16.5 and P0, in heart, liver, forebrain, midbrain and hindbrain. EpiTensor performed tensor analysis in the 4 dimensional space composed of tissue, time point, histone mark, and locus dimensions. The peaks in the eigenlocus vectors represent co-variation of histone modification signals that indicate 3D interactions of the corresponding loci. We considered the first 40 eigenlocus vectors, capturing on average 96.98% of total variance. The peaks in each eigenlocus vector were called by comparing to randomly shuffled background. FDR and p-value were calculated for each peak. In this study we used a stringent FDR=0.005 as cutoff.

### Lineage tree construction

Lineage trees were constructed by running the FastME algorithm^25^ on normalized ranking matrix, which was obtained from row-wise z-score transformation of the original matrix.

### Identification of housekeeping TFs and driver TFs for tissues

The output of Taiji pipeline is a matrix, consisting of TFs’ ranking scores from different experiments. The rows and columns of the matrix represent TFs and experiments respectively. To identify housekeeping TFs, we first selected TFs with average ranking scores larger than 3e-3, corresponding to the top 10% of all TFs. Next, we removed TFs of which the coefficient of variance (CV) are greater than 0.5, keeping the rest as the housekeeping TFs. To identify driver TFs, we first removed the rows (TFs) with CVs less than 1. We then grouped the columns (experiments) from various stages together if they are from the same tissue, and averaged the ranking scores in each group. Assuming the average ranking scores are normally distributed, we calculated the deviation from the center for each score and computed the p-value. A 0.01 p-value cutoff was used for calling driver TFs.

## Code Availability

Taiji software is available at http://wanglab.ucsd.edu/star/taiji.

## Acknowledgement

This project is partially supported by NIH (U54HG006997 and R01HG009626) and CIRM (RB5 07012).

## Author Contributions

K.Z. and W.W. conceived the study and interpreted the results; K.Z. designed the Taiji software; K.Z. and M.W. performed the data analysis with contributions from Y.Z.; K.Z. and W.W. wrote the manuscript with contributions from M.W. and Y.Z.; W.W. supervised the project.

---

**Extended Data Figure 1:**
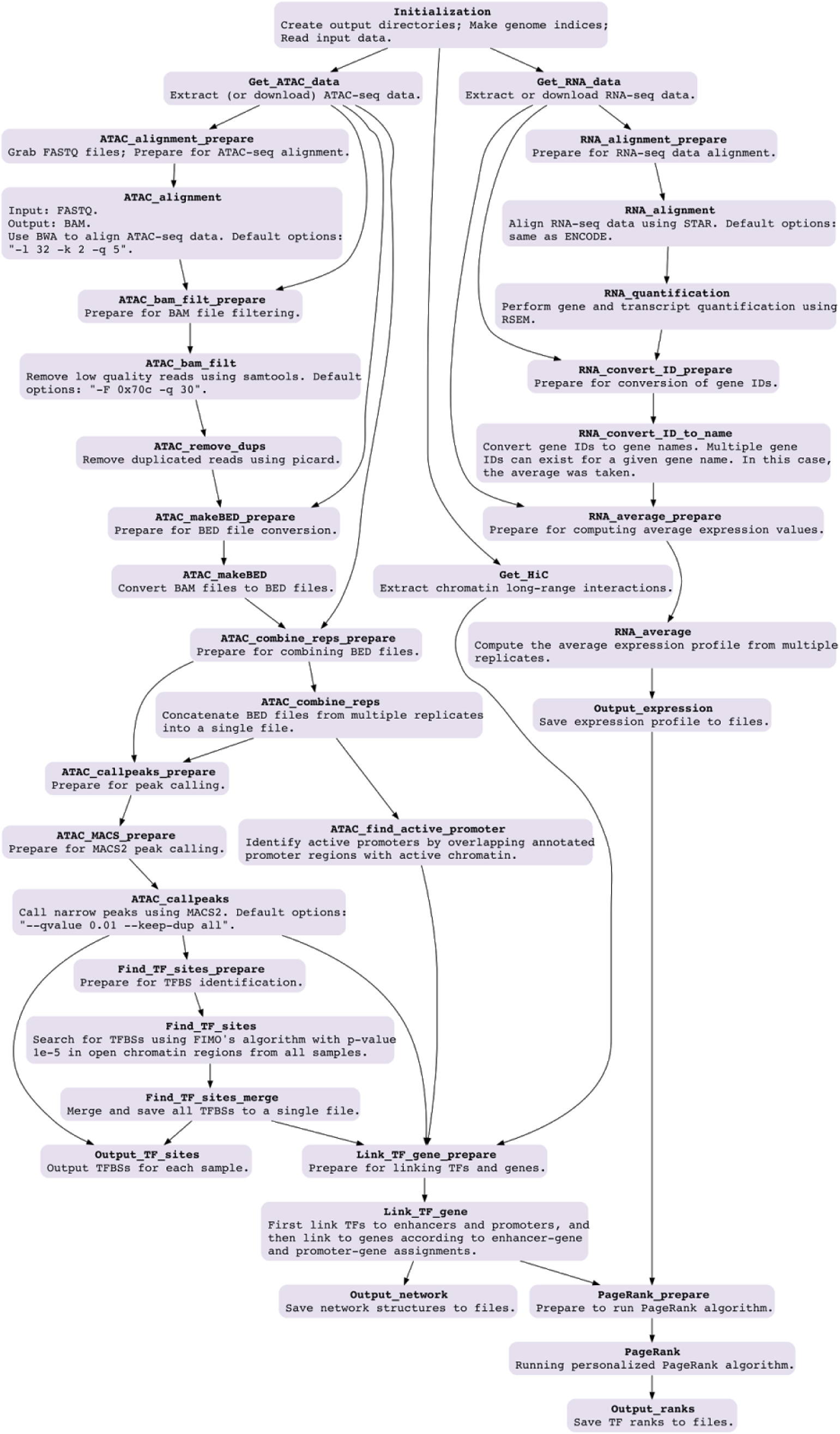
The complete design of Taiji pipeline. Each box represents a computation step.

**Extended Data Figure 2:**
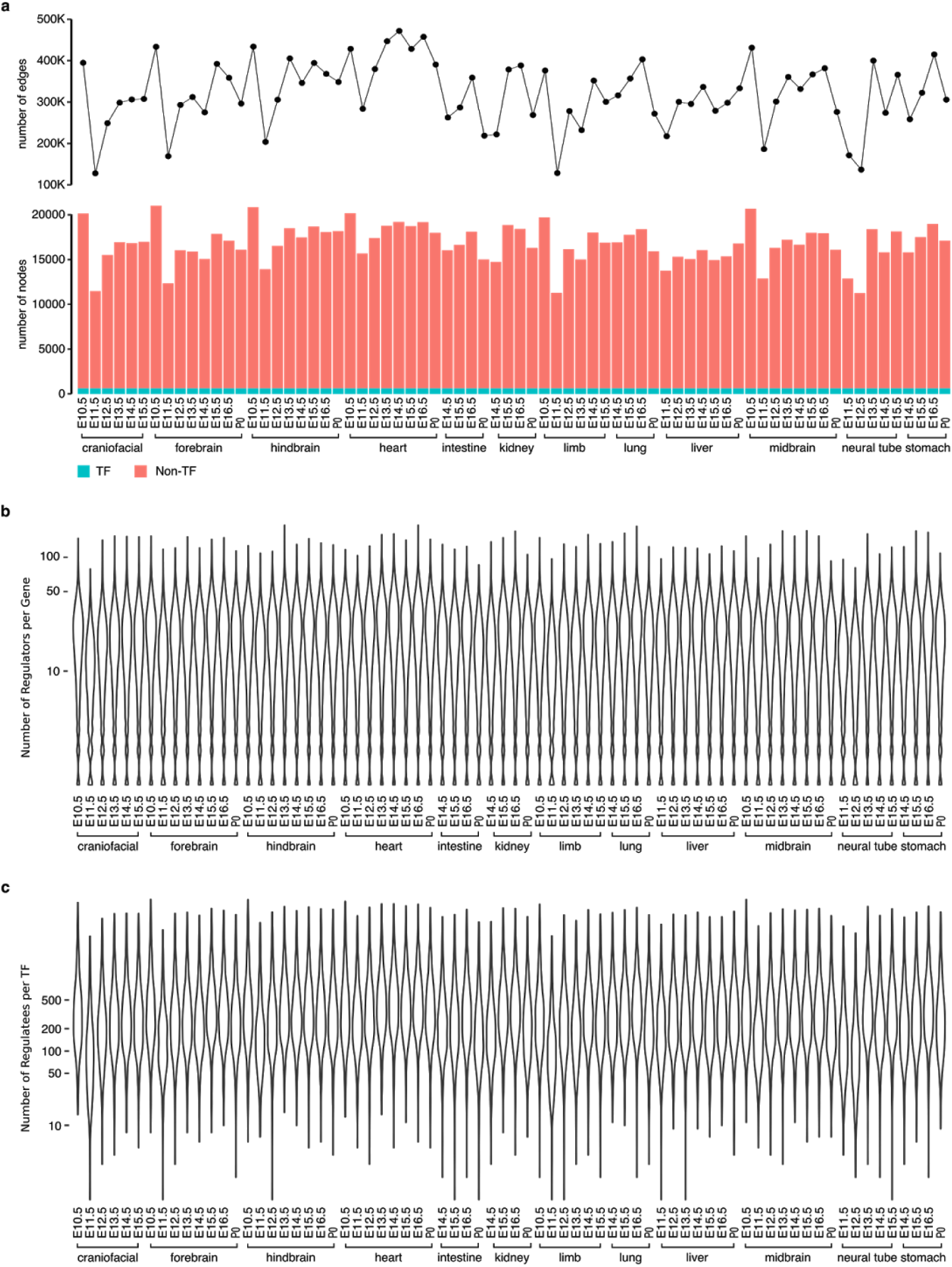
The topological properties of genetic networks. **a**, The number of edges (top) and nodes (bottom) in each genetic network; **b**, The distribution of the number of regulators per gene for each genetic network; **c**, The distribution of the number of regulatees per TF for each genetic network.

**Extended Data Figure 3:**
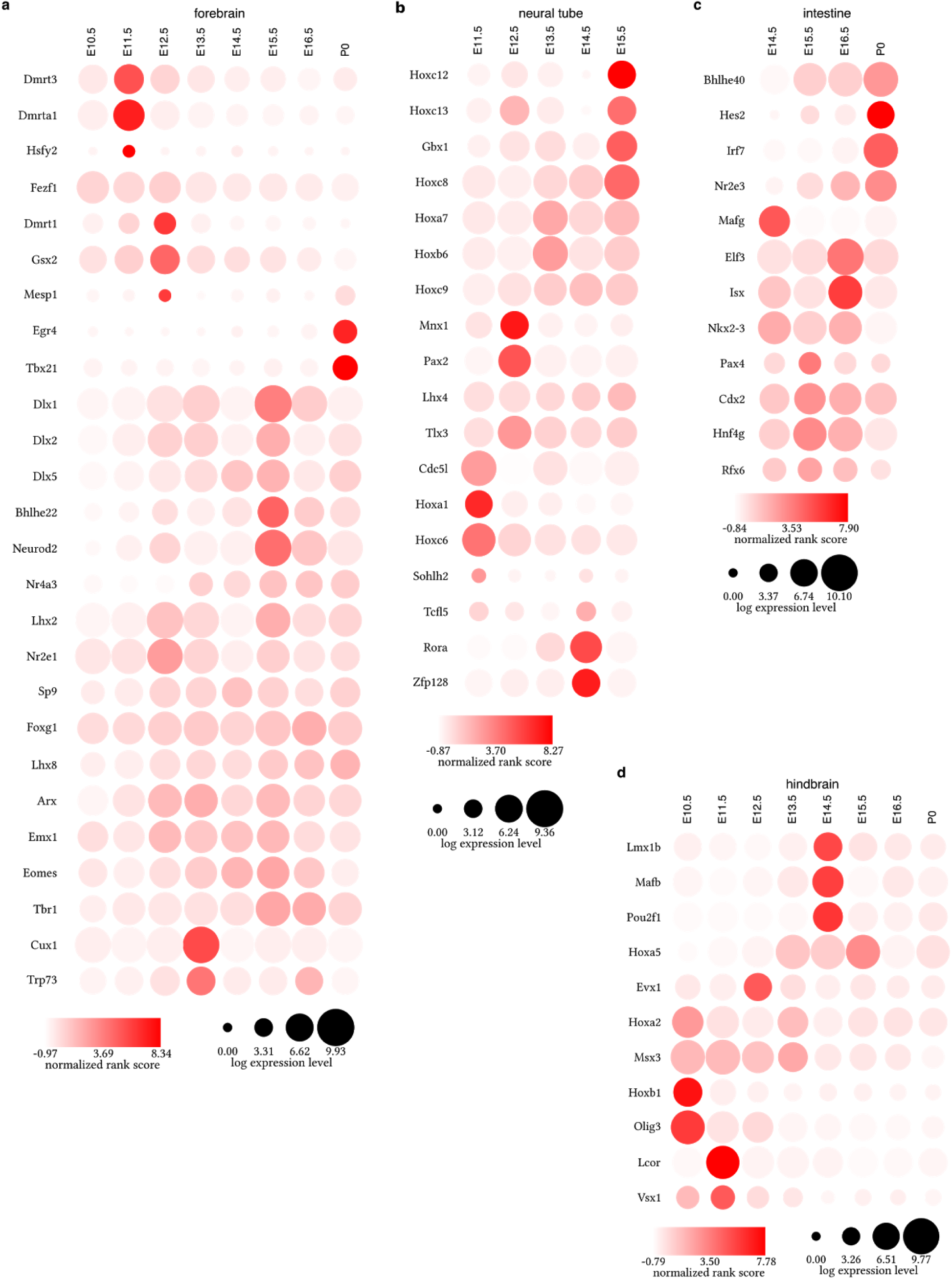
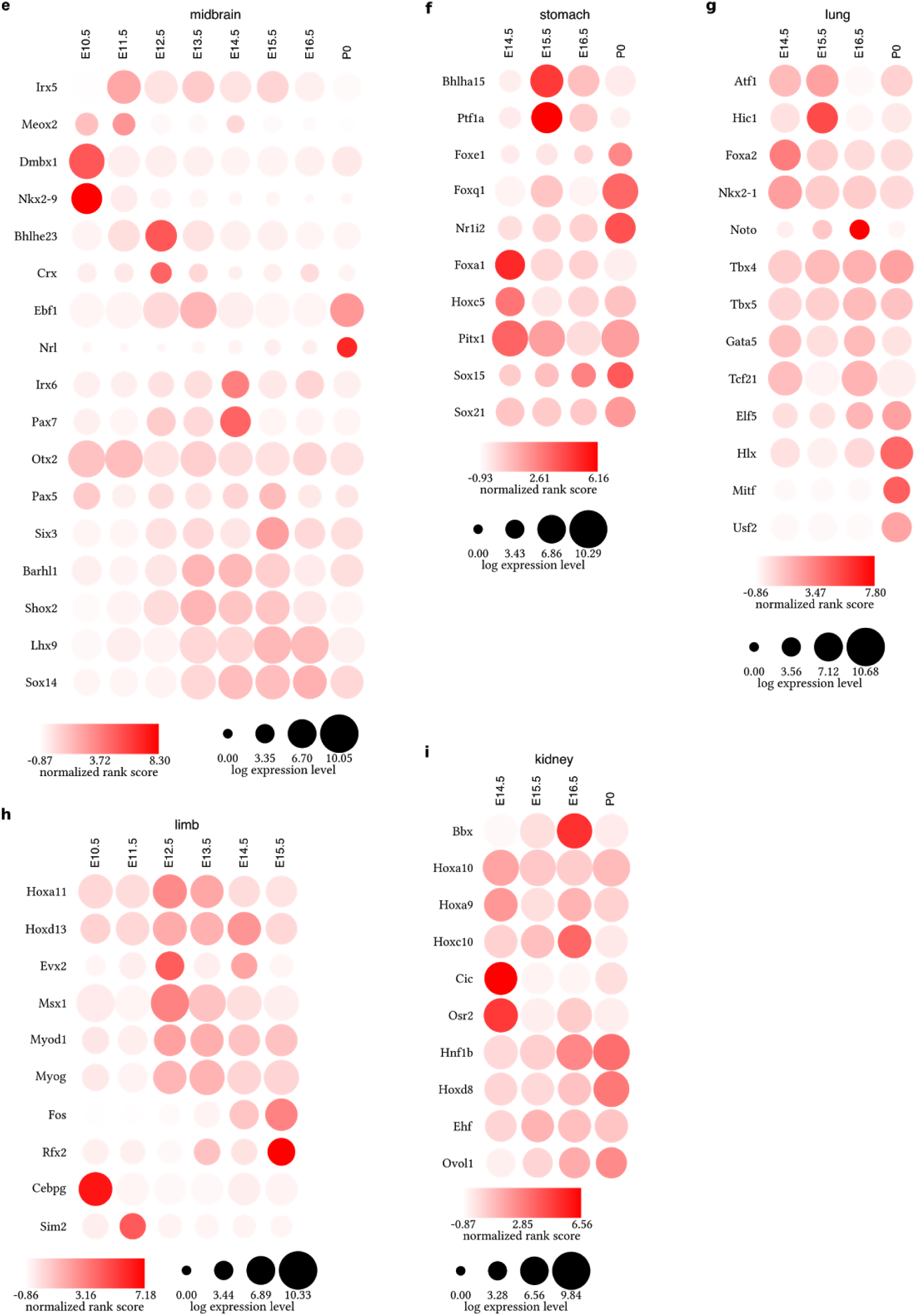
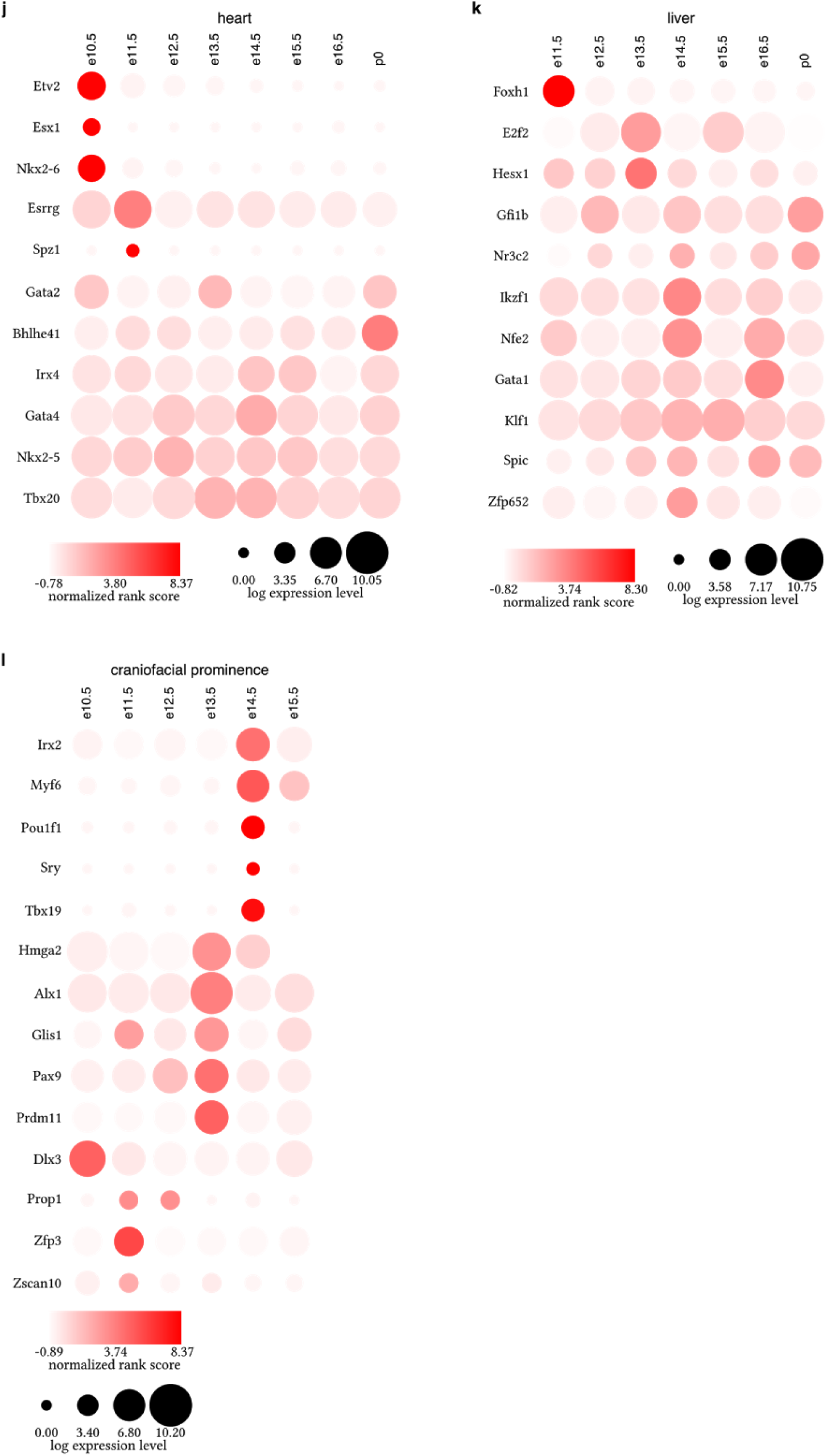
Identification of driver TFs in 12 tissues. **a**, forebrain. **b**, neural tube. **c**, intestine. **d**, hindbrain. **e**, midbrain. **f**, stomach. **g**, lung. **h**, limb. **i**, kidney. **j**, heart. **k**, liver. l, craniofacial prominence.

**Extended Data Figure 4:**
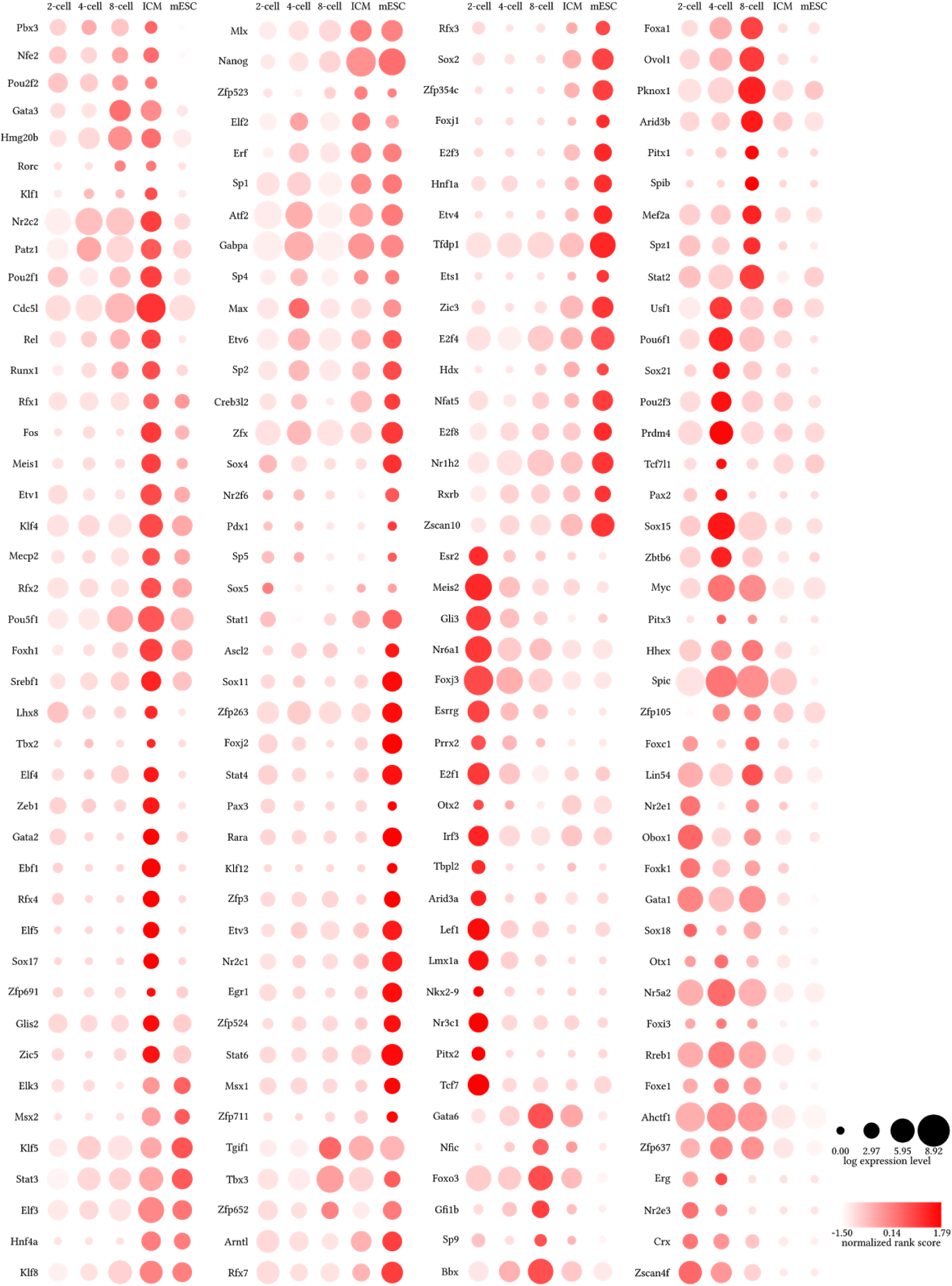
Identification of driver TFs in early mouse embryonic development, including 2-cell, 4-cell, 8-cell stages, ICM and ESC.

**Extended Data Figure 5:**
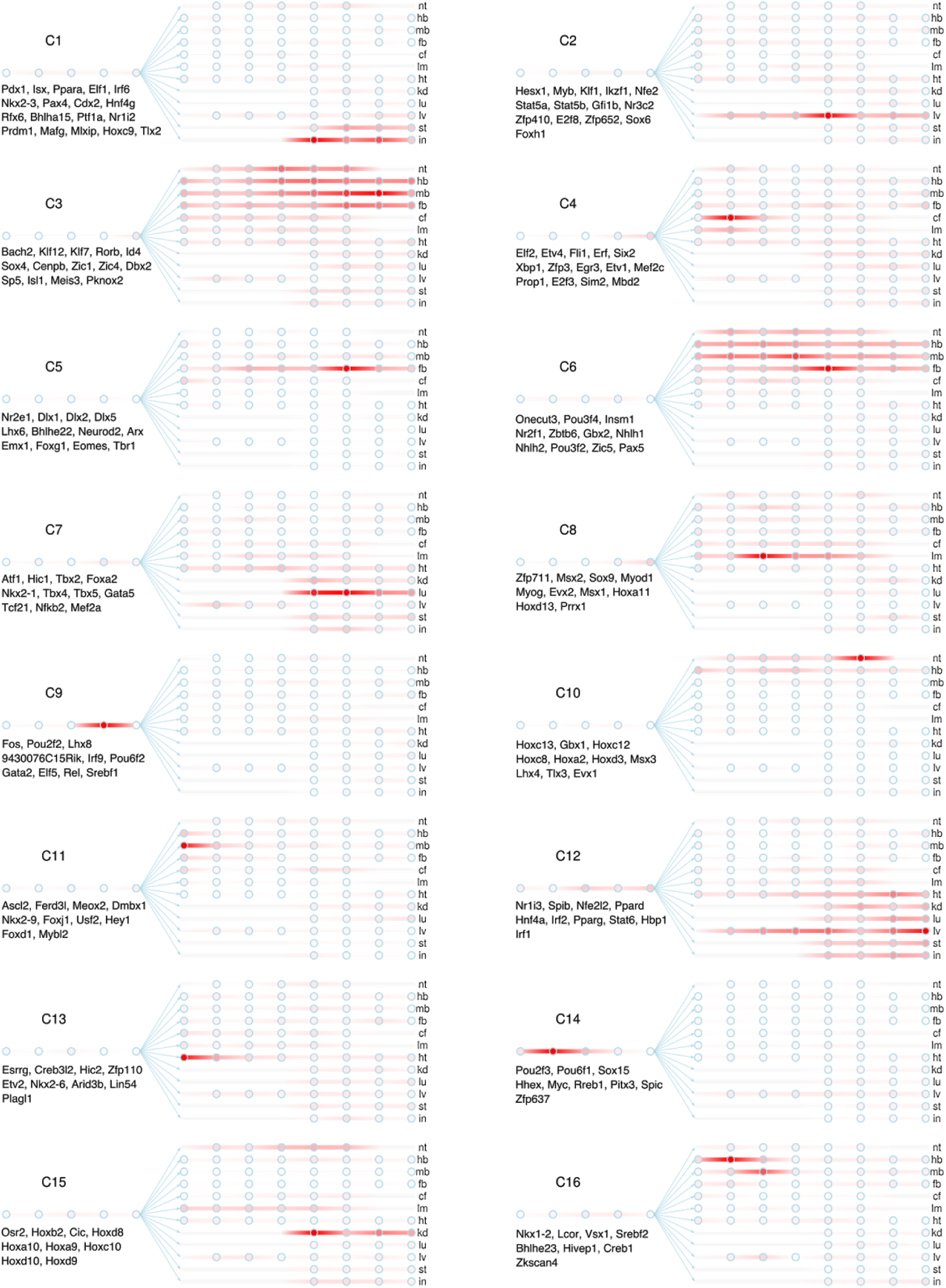
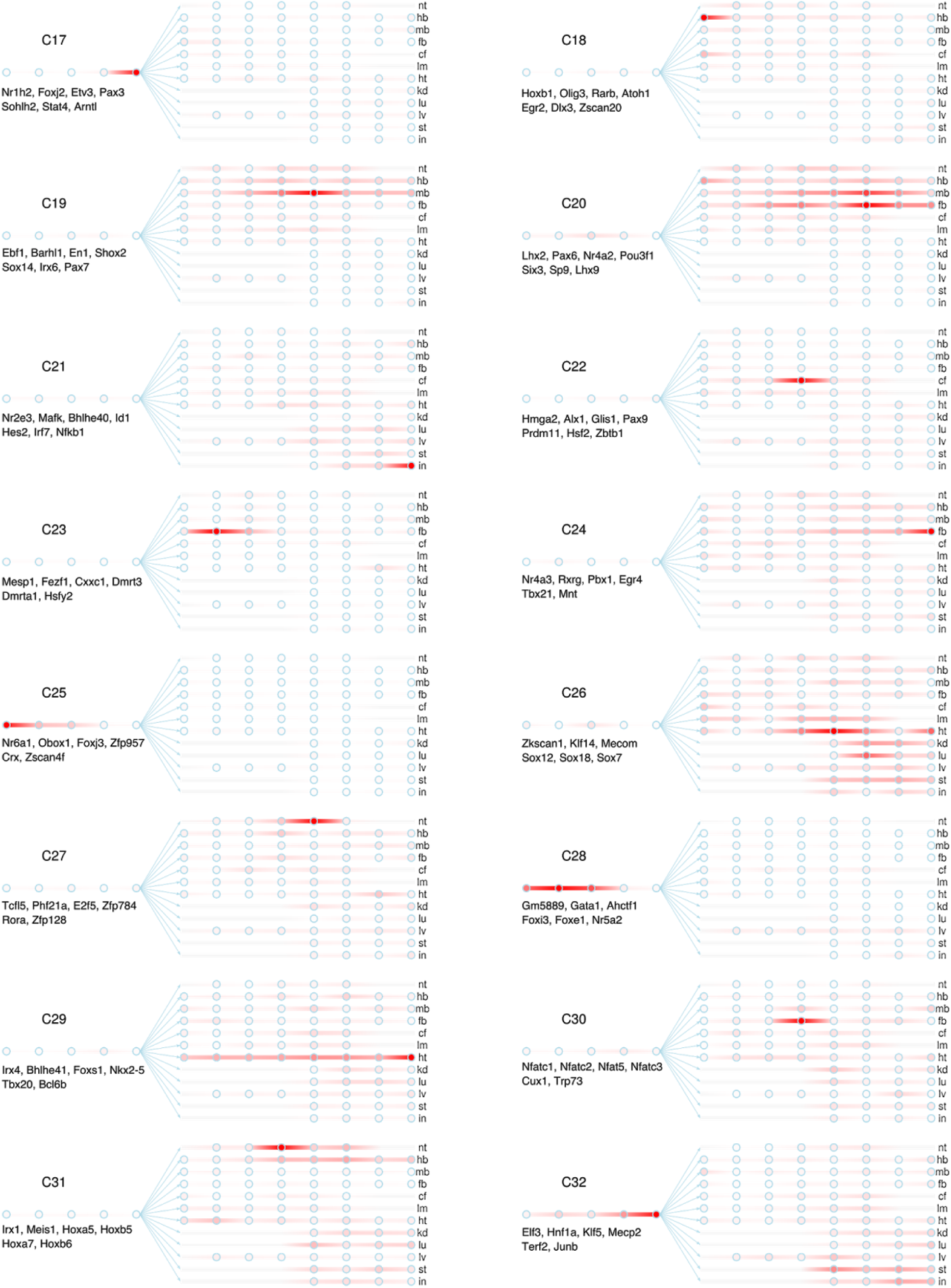
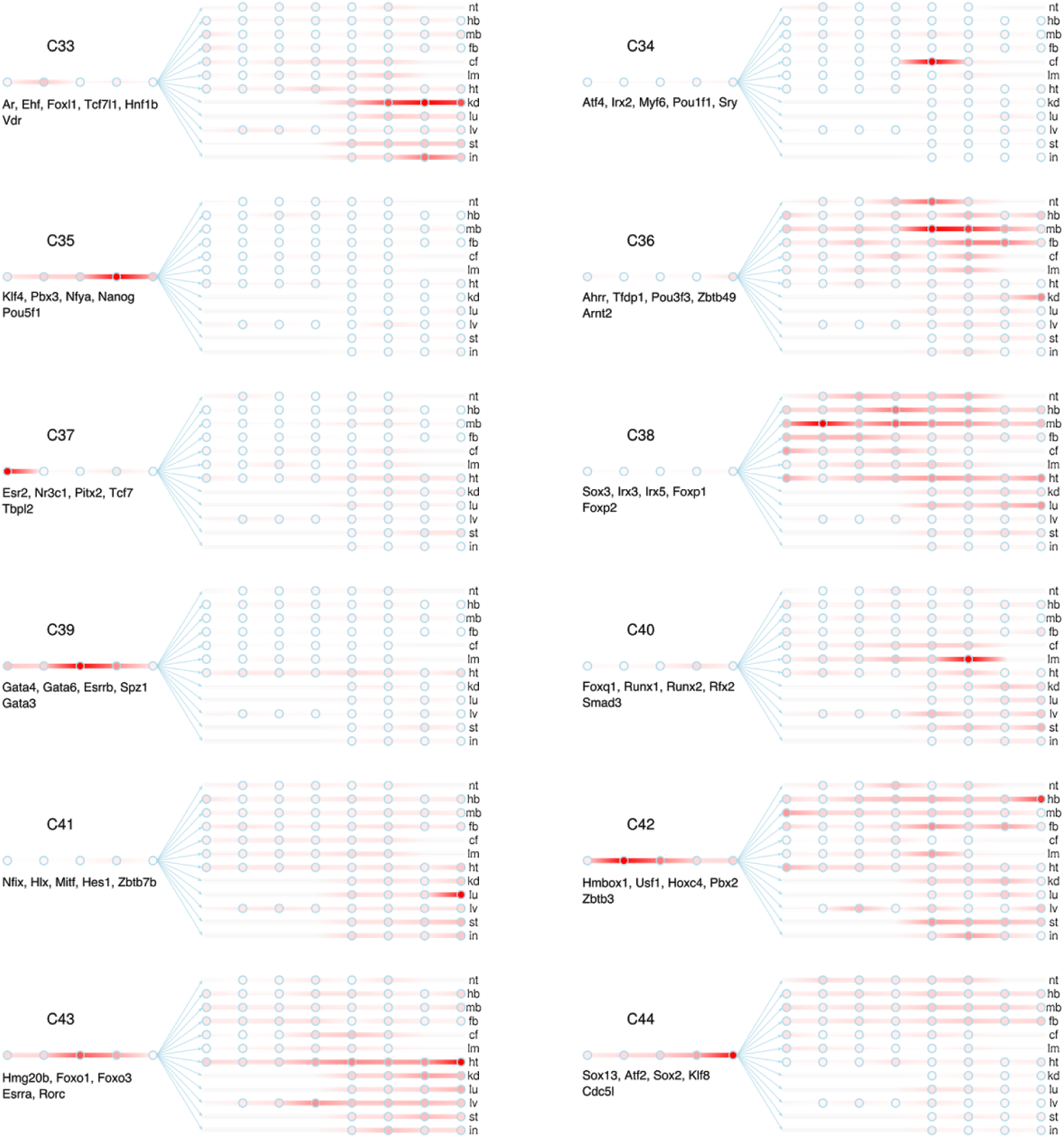
Transcriptional waves direct tissue differentiation during mouse embryogenesis. A total of 44 dynamic patterns of ranking scores were identified using hierarchical clustering algorithm. Circles in each panel represent developmental stages. From left to right, they are 2-cell, 4-cell, 8-cell, ICM, ESC, E10.5, E11.5, E12.5, E13.5, E14.5, E15.5, E16.5 and P0 respectively. TF members are shown below the names of clusters.

